# Targeting of human mitochondrial DNA with programmable pAgo nuclease

**DOI:** 10.1101/2025.11.20.689640

**Authors:** Beatrisa Rimskaya, Ekaterina Kropocheva, Elza Shchukina, Egor Ulashchik, Daria Gelfenbein, Lidiya Lisitskaya, Vadim Shmanai, Svetlana Smirnikhina, Andrey Kulbachinskiy, Ilya Mazunin

## Abstract

Manipulating the mitochondrial genome remains a major challenge in genetic engineering, largely due to the mitochondrial double-membrane structure. While recent advances have expanded the genetic toolkit for nuclear and cytoplasmic targets, precise editing of mtDNA has remained elusive. Here we report the first successful mitochondrial import of a functional RNA-guided prokaryotic Argonaute protein from the mesophilic bacterium *Alteromonas macleodii* (AmAgo). By directing AmAgo to the single-stranded D- or R-loop region of mtDNA using synthetic RNA guides, we observed a nearly threefold reduction in mtDNA copy number in human cell lines. This proof of concept study demonstrates that a bacterial Argonaute can remain active within the mitochondrial environment and influence mtDNA levels. These findings lay the groundwork for further development of programmable systems for mitochondrial genome manipulation.

**Highlights:** AmAgo targeted to mitochondria significantly reduces human mtDNA copy number Mitochondrial localization of AmAgo is essential for changes in the mtDNA content AmAgo effects on mtDNA are modulated by exogenous guide RNAs

*Graphical abstract:* 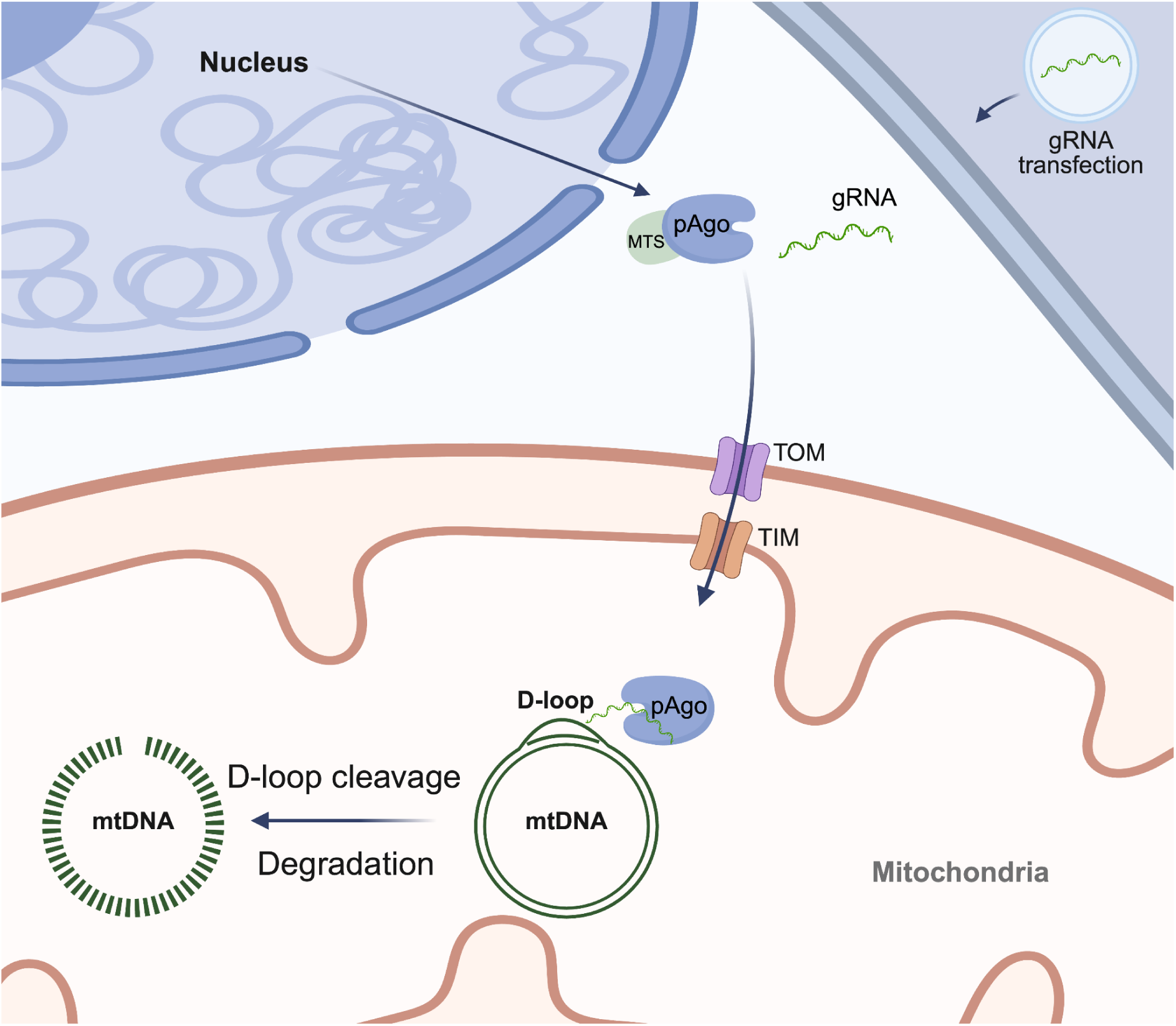

## Introduction

The mitochondrial genome (mtDNA) forms an independent genetic system distinct from the nucleus, encoding core components of the oxidative phosphorylation machinery as well as key tRNAs and rRNAs. In addition to ATP synthesis, mitochondria participate in essential cellular processes including calcium signaling, apoptosis, heme biosynthesis, and amino acid metabolism.^1^ Mutations in mtDNA can disrupt these pathways and lead to a wide spectrum of disorders collectively known as mitochondrial diseases,^2^ which range from early-onset neurodegenerative syndromes to multisystemic metabolic conditions.

Despite recent progress, precise manipulation of mtDNA remains technically challenging. Current tools - including modified transcription activator-like effector nucleases (TALEN), zinc-finger nucleases, restriction endonucleases, artificial meganucleases, and CRISPR/Cas systems - have been adapted to target the mitochondrial genome, primarily for shifting heteroplasmy or correcting transition mutations.^3^ However, these tools exhibit significantly lower efficiency compared to their nuclear genome counterparts due to substantial challenges associated with delivering nucleases and guiding nucleic acids into mitochondria. Moreover, broader manipulations, such as transversion editing, locus deletion, or sequence integration within mtDNA, remain technically unresolved. Thus, novel methodological advancements are urgently required.

Prokaryotic Argonaute proteins (pAgos) offer a promising alternative to Cas nucleases.^4,5^ In contrast to eukaryotic Argonautes, which play a central role in RNA interference and act on RNA targets, most prokaryotic Argonautes recognize DNA substrates.^6,7^ Furthermore, compared to CRISPR-associated nucleases such as Cas9 (∼160 kDa) and Cas12a (∼150 kDa), Argonautes are considerably smaller (e.g., AmAgo, ∼65 kDa^8^), a property that may facilitate their mitochondrial import. Cas nucleases typically rely on guide RNAs more than 40 nucleotides in length and require the presence of protospacer-adjacent motifs (PAMs) flanking the target site, which restricts targeting flexibility.^9^ In contrast, pAgos use short single-stranded RNA or DNA guides (typically 16–20 nt) and are PAM-independent, enabling programmable cleavage across a broader range of DNA targets.^5,7,10,11,12,13,14^

First attempts to use the Argonaute from *Natronobacterium gregoryi* (NgAgo) with guide DNA for editing of genomic DNA in mammalian cells were found to be irreproducible.^15,16,17,18^ At the same time, it was reported that NgAgo was able to inhibit gene expression in eukaryotic cells in the absence of DNA editing.^19,20^ Following attempts to use NgAgo and TtAgo from *Thermus thermophilus* for programmable DNA binding were unsuccessful.^21^

Among pAgos, the Argonaute from *Alteromonas macleodii* (AmAgo), used in this study, utilizes single-stranded RNA guides to catalyze site-specific cleavage of single-stranded DNA (ssDNA).^8^ This biochemical property makes AmAgo particularly suitable for targeting the mitochondrial D-loop and R-loop, naturally exposed ssDNA structures that play a central role in replication and transcription initiation. Furthermore, the high copy number of mitochondrial genomes (hundreds to thousands per cell) may enhance the likelihood of Argonaute-mediated recognition and cleavage of the target loci.

However, pAgos have not previously been demonstrated to function in mitochondria and there were no attempts to use pAgos guided by RNA as a programmable DNA-targeting system in eukaryotic cells. Their application has been limited by uncertainty around mitochondrial import, guide delivery, and enzymatic activity in mammalian systems. Here, we report the first successful delivery of AmAgo into human mitochondria. Targeting the D-loop and R-loop regions with synthetic RNA guides led to a reproducible ∼3-fold reduction in the mtDNA copy number. These results establish a foundation for developing compact, programmable, and PAM-independent tools for mitochondrial genome engineering.

## Materials and Methods

### Molecular cloning

All transfer plasmids were generated by modifying the lentiviral pUltra vector (Addgene plasmid #24129) using standard molecular biology techniques. Constructs encoding mitochondria-targeted prokaryotic Argonaute (pAgo) were assembled by PCR amplification of the AmAgo coding sequence, a 3×FLAG tag, and a mitochondrial targeting sequence (MTS) derived from the subunit 9 of *Neurospora crassa* mitochondrial ATP synthase (Su9), selected based on previously published data.^22^

To generate lentiviral transfer plasmids, the GFP expression cassette in pUltra was replaced with sequences encoding either 3×FLAG-MTS or 3×FLAG alone. These inserts were cloned into the vector using *Nhe*I and *Asi*GI restriction sites. The AmAgo coding region, amplified from a separate plasmid template, was subsequently inserted into the construct via *Xho*I and *Nhe*I sites. Primer sequences used for template amplification are listed in Supplementary Table S1.

### Protein expression and purification

For expression of wild-type and mutant AmAgo proteins, *E. coli* BL21(DE3) cells carrying the pET28-based expression plasmid^8^ were grown in the LB medium with 50 µg/ml kanamycin at 37 °C until OD_600_ 0.35, cooled down to 16°C, induced by the addition of 0.1 mM IPTG and grown for 16 h at 16°C. The cells were collected by centrifugation and stored at -80 °C.

To purify AmAgo, the cells were resuspended in 50 ml of buffer A1 (40 mM Tris-HCl pH 7.4, 500 mM NaCl, 0.1 mM DTT, 5% glycerol, 10 µg/mL PMSF) and disrupted using an ultrasonic homogenizer (QSonica Q125). The lysate was cleared by centrifugation at 35,000 g for 30 min, supplemented with 10 mM imidazole and loaded onto a HisTrap Fast Flow crude column (GE Healthcare). The column was washed with the same buffer without PMSF containing 30 mM imidazole and then AmAgo was eluted with 5 ml of the buffer containing 300 mM imidazole. Fractions containing AmAgo were combined, diluted with 4 volumes of buffer B1 (40 mM Tris-HCl pH 7.4, 0.5 mM EDTA, 0.1 mM DTT, 5% glycerol) and loaded onto a HiTrap SP HP cation-exchange column equilibrated with buffer B1 supplemented with 100 mM NaCl. AmAgo was eluted with a linear NaCl gradient. Samples containing AmAgo were concentrated by ultrafiltration using Amicon-30K.

### Analysis of nucleic acid cleavage by AmAgo in vitro

Guide RNAs and target DNAs were synthesized on an ASM-2000 automated DNA/RNA synthesizer (Biosset Ltd.) at a 0.5 μmol scale using 1000 Å Universal CPG solid support (Primetech ALC, loading capacity: 41 μmol/g) under conditions similar to those previously described^23^. The sequences of all oligonucleotides are listed in Supplementary Table S2.

The nuclease activity of AmAgo was tested *in vitro* using synthetic oligonucleotides in a reaction buffer containing 10 mM Tris-HCl pH 7.9, 100 mM NaCl, 5 mM MgCl_2_, and 5% glycerol at 37°C. AmAgo (500 nM final concentration) was mixed with guide RNA oligonucleotide (200 nM), incubated for 15 min at 37 °C for guide loading, then target DNA or RNA was added (100 nM), and the reaction was stopped after indicated time points by adding an equal volume of a stop-solution containing 8 M urea, 1 mM EDTA, 0.005% Bromophenol Blue. The reaction products were separated by 19% denaturing urea PAGE. The gels were stained with SYBR Gold (Invitrogen) and visualized with a Typhoon FLA 9500 scanner (GE Healthcare).

### Cell culture

Human embryonic kidney 293T (HEK293T) cells and fibroblasts from healthy donor were maintained in high-glucose Dulbecco’s Modified Eagle Medium (DMEM) supplemented with 10% bovine calf serum, 1× antibiotic-antimycotic solution (final concentrations: 100 U/mL penicillin, 100 μg/mL streptomycin, and 0.25 μg/mL amphotericin B), 1× sodium pyruvate, 1× non-essential amino acids, and 1× GlutaMAX. Cells were cultured in T75 flasks (75 cm² surface area) under standard conditions (37°C, 5% CO_2_).

### Guide RNA transfection

One day prior to transfection, cells were seeded into 48-well plates pre-coated with gelatin at a density of 70,000 cells per well to achieve approximately 50% confluency on the day of transfection. Transfection was carried out using Lipofectamine 2000 (Thermo Fisher Scientific) following the manufacturer’s protocol (https://www.thermofisher.com/ru/ru/home/references/protocols/cell-culture/transfection-prot ocol/lipofectamine-2000.html). For each well, two transfection mixes were prepared. Mix A contained 500 ng of guide RNA in 25 µL of Opti-MEM Reduced Serum Medium (Thermo Fisher Scientific). Mix B consisted of 1 µL of Lipofectamine 2000 diluted in 25 of Opti-MEM. The two mixtures were combined and incubated at room temperature for 20 minutes to allow complex formation. Subsequently, 50 µL of the resulting transfection complex was added to each well. Cells were incubated under standard conditions for 48 hours post-transfection. Total DNA was then extracted from each well for downstream analysis.

### Lentivirus particle assembly

HEK293T cells were seeded in 6-well plates at a density of 1 × 10^6^ cells per well to reach approximately 80% confluency after 24 hours. Two hours prior to transfection, the culture medium was replaced with serum-free Opti-MEM (Thermo Fisher Scientific). For lentiviral transfection, the lentiviral transfer vector, psPAX2 packaging plasmid (Addgene plasmid #12260), and pVSV-G envelope plasmid (Addgene plasmid #138479) were mixed at equimolar ratios. A total of 5 μg of plasmid DNA was diluted in 80 μL of distilled water, followed by the addition of 10 μL of 2M CaCl_2_. The DNA mixture was aerated vigorously using a sterile 200 μL pipette tip. While bubbling, 100 μL of 2× HEPES-buffered saline (280 mM NaCl, 50 mM HEPES, 1.5 mM Na_2_HPO_4_, pH 7.0) was added dropwise to initiate calcium phosphate-DNA precipitate formation. The solution was briefly vortexed and incubated at room temperature for 15 minutes. After a second gentle vortex, the transfection mixture was added dropwise to the cells. Plates were gently rocked to distribute the precipitate evenly and then returned to a 37°C CO_2_ incubator.

Four to six hours post-transfection, the medium was replaced with a fresh growth medium supplemented with 10% heat-inactivated fetal bovine serum (FBS). At 16 hours post-transfection, sodium butyrate was added to the culture medium at a final concentration of 2 mM to enhance viral production. Virus-containing supernatants were collected 48 hours later. Conditioned media were clarified by centrifugation at 1500 rpm for 5 minutes at 4°C and then passed through 0.45 μm PVDF syringe filters (Millex-HV, Millipore) to remove cellular debris. The resulting viral preparations were aliquoted and stored at –80°C until use.

### Lentiviral transduction

Lentiviral transduction was performed by adding viral particles to HEK293T cells cultured in DMEM supplemented with 10% FBS and 8 μg/mL polybrene. The virus was added at a multiplicity of infection (MOI) of 5, which has been previously reported as optimal for the HEK293T cell line (https://cdn.origene.com/assets/documents/lentiviral/recommended_lentivirus_moi_for_com mon_cell_lines.pdf). After 6 hours of incubation, the viral medium was removed and replaced with fresh culture medium. Forty-eight hours post-transduction, cells were analyzed by flow cytometry to quantify GFP-positive cells as a measure of functional viral titer. A control condition corresponding to a multiplicity of infection of five was established based on GFP-positivity and calculated following the Cellecta Lentiviral Titer Calculation Manual (https://manuals.cellecta.com/lentiviral-construct-packaging-and-transduction/v10a/en/topic/l entiviral-titer-calculation). This GFP-positive sample was subsequently used as a reference to evaluate the multiplicity of infection in experimental conditions by quantitative real-time PCR (qPCR).

### MtDNA copy number evaluation

Total DNA was extracted from HEK293T cells using the ExtractDNA Blood & Cells Kit (Evrogen, Russia) according to the manufacturer’s protocol. MtDNA copy number was quantified by real-time PCR using primers targeting the mitochondrial D-loop region or MT-ND1 gene and the single-copy nuclear gene Beta-2-Microglobulin (B2M) as a reference for nuclear DNA (nDNA). Each 25 μL reaction contained 1× SYBR Green I PCR mix (Evrogen, Russia), 300 nM of each primer, and 50 ng of total genomic DNA. Amplification was performed using a QuantStudio 5 system (Applied Biosystems, USA) under standard cycling conditions.

For each condition, three independent biological replicates were analyzed, each with three technical replicates. The relative mtDNA copy number was calculated using the 2^−ΔΔCt^ method, comparing Ct values of D-loop and B2M, and normalized to control samples.

### Immunocytochemical assay for AmAgo

Fibroblasts were seeded onto gelatin-coated 8-well chamber slides at a density of 3.0 × 10^5^ cells per well, 48 hours after transduction. After 24 hours of incubation, cells were washed once with phosphate-buffered saline (PBS) and stained with 100 nM MitoTracker CMX-ROS (Thermo Fisher Scientific) for 30 minutes at 37°C in a 5% CO_2_ incubator to visualize mitochondria. Following staining, cells were washed twice with PBS, fixed with 4% paraformaldehyde for 15 minutes at room temperature, and permeabilized with 0.5% Triton X-100 in PBS for 10 minutes. Non-specific binding was blocked with 4% bovine serum albumin (BSA) in PBS for 1 hour at room temperature. Cells were then incubated overnight at 4 °C with a mouse anti-FLAG primary antibody (Sigma, Monoclonal ANTI-FLAG M2; 1:500 dilution in PBS containing 4% BSA). After primary antibody incubation, cells were washed three times with PBS and incubated for 1 hour at room temperature with a goat anti-mouse Alexa Fluor 488-conjugated secondary antibody (Thermo Fisher Scientific; 1:400 dilution in PBS). Slides were washed extensively with PBS and mounted using an antifade mounting medium. Fluorescence imaging was performed using a fluorescence microscope equipped with appropriate filters (Agilent BioTek Lionheart FX). Microscopy images were processed and analyzed using ImageJ software. Co-localization was assessed by extracting fluorescence intensity profiles along linear regions of interest (ROIs) to evaluate the spatial overlap between MitoTracker and FLAG signals.

### In silico selection of MTS

Physicochemical parameters relevant to mitochondrial targeting - net charge, isoelectric point (pI), and grand average of hydropathy (GRAVY) - were calculated using the ExPASy ProtParam tool (https://web.expasy.org/protparam/) for each MTS alone and for the full-length fusion proteins (Supplementary Table S3).

### Flow cytometry

Cells were collected 48 hours post-transduction, dissociated by pipetting, and resuspended in PBS without calcium and magnesium. GFP fluorescence was measured in the FITC-A channel using a CytoFLEX S V2-B2-Y2-R0 flow cytometer (Beckman Coulter, Brea, CA, USA). Data were analyzed in CytExpert v2.4. GFP-positive gates were defined relative to non-transduced controls. At least 10,000 events were acquired per sample.

### Statistical analysis

All datasets represent three independent experiments, each including ≥3 technical replicates. Technical replicates were averaged per biological replicate. Statistical significance was assessed using two-tailed *t*-tests in Python (v3.10.9); paired or unpaired tests were applied depending on the experimental design. Data are plotted as mean ± standard deviation (SD). Statistical significance is represented as follows: ns, not significant (P > 0.05); *P ≤ 0.05; **P < 0.01.

## Results

### Import of the AmAgo nuclease into human mitochondria

To facilitate mitochondrial import of AmAgo, we evaluated four candidate mitochondrial targeting sequences (MTSs) *in silico*. Sequences derived from cytochrome c oxidase subunit 8A (COX8A), ATPase subunit 9 (Su9) from *Neurospora crassa*, a cryptic MTS from the mammalian autophagy-related protease ATG4D, and mitochondrial superoxide dismutase 2 (SOD2) were fused *in silico* to the N-terminus of AmAgo via a 3xFLAG tag, as previously described.^22^ All constructs displayed negative GRAVY scores, consistent with hydrophilic properties and expected solubility. Since mitochondrial import is initiated by recognition of positively charged amphipathic helices at the N-terminus by the TOM complex, and driven in part by the negative membrane potential (Δψ) across the inner membrane, local electrostatic potential at the MTS is considered a key determinant of import efficiency.^24^ In contrast, total charge across the full-length fusion may be less predictive of targeting performance. *In silico* analysis of the MTS-AmAgo fusions predicted that all constructs retained hydrophilic properties and solubility. Su9 and ATG4D retained strong net positive charges in the MTS region (+13 and +10, respectively), whereas the SOD2 construct showed the lowest MTS charge (+3) and was the only one with a negative net charge (–2) in the full-length context. Notably, although the ATG4D fusion had a higher overall charge (+12), Su9 exhibited the strongest net positive charge specifically at the N-terminus. Based on these parameters, Su9 was prioritized for experimental testing.

A lentiviral vector encoding Su9-3xFLAG-AmAgo or 3xFLAG-AmAgo was introduced into the fibroblasts for further immunocytochemistry analysis (Figure 1A). Fluorescence microscopy revealed robust expression of the Su9-tagged AmAgo and its clear colocalization with mitochondrial markers (Figure 1B). To quantify colocalization, fluorescence intensity profiles were extracted along shared linear region of interest (ROI) using ImageJ (Figure 1C). Pearson’s correlation coefficient calculated from raw pixel values yielded R = 0.94, indicating robust mitochondrial localization. In contrast, AmAgo lacking an MTS showed diffuse cytoplasmic and nuclear distribution (Figure 1B). These results confirm that Su9 is an effective and biophysically favorable MTS for the targeted delivery of programmable nucleases such as AmAgo into the mitochondrial matrix.

**Figure 1.**
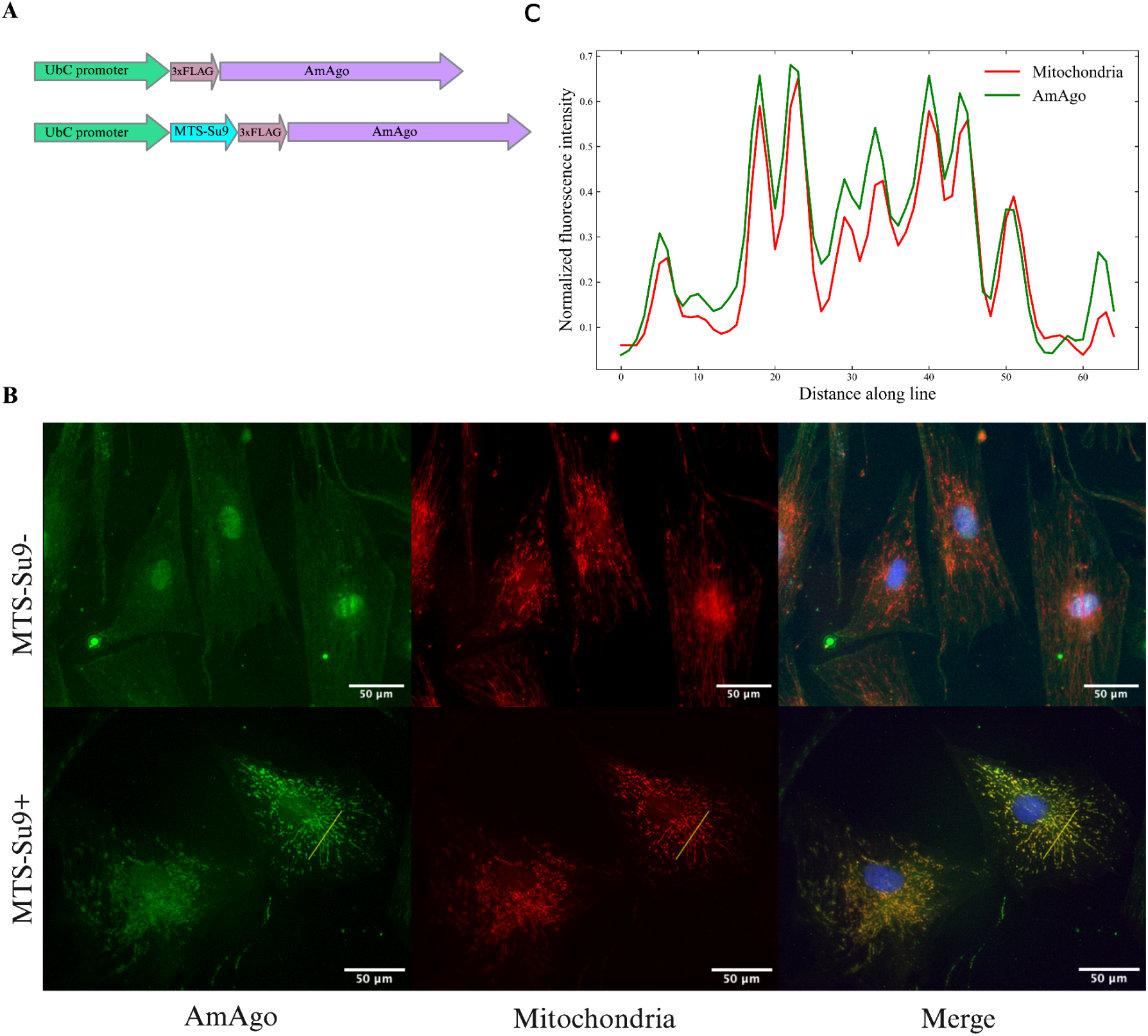
Analysis of AmAgo colocalization in mitochondria. **A** – Schematic representation of expression constructs encoding the prokaryotic Argonaute protein AmAgo under the control of the *UbC* promoter. In the experimental construct (bottom), AmAgo is fused to the mitochondrial targeting sequence (MTS) Su9 at the N-terminus and tagged with a 3×FLAG epitope. The control construct (top) lacks the MTS sequence. **B** – **Immunofluorescence analysis of AmAgo localization in human fibroblasts.** Fibroblasts were used instead of HEK293T cells to improve visualization of the mitochondrial network. Cells were stained with anti-FLAG antibody (green) to detect AmAgo, MitoTracker CMXRos (red) to label mitochondria, and DAPI (blue) for nuclear staining. In cells expressing MTS-Su9–AmAgo (bottom row), a clear overlap between AmAgo and mitochondria is observed in the merged image. In contrast, cells expressing MTS-lacking AmAgo (top row) show diffuse distribution of the protein throughout the cytoplasm and nucleus. **С** – **Fluorescence intensity profile analysis along a selected ROI.** Normalized fluorescence intensities of AmAgo (green) and mitochondria (MitoTracker, red) were plotted along a yellow line drawn across a representative cell to assess signal colocalization. Pearson’s correlation coefficient calculated from raw fluorescence intensity profiles confirmed strong colocalization (R = 0.94).

### Guide RNA design and in vitro testing

We previously demonstrated that AmAgo directs cleavage of ssDNA targets using unphosphorylated RNA guides containing a 5’-adenine.^8^ To evaluate its functional activity in a mitochondrial context, we designed RNA guides to target the human mtDNA control region (particularly, the D-loop and R-loop), a regulatory region with a triple-stranded structure composed of a DNA duplex with an invading DNA strand and an exposed single-stranded loop (Figure 2A, Supplementary Table S2). This region plays a central role in mtDNA replication and transcription initiation and offers structurally accessible single-stranded segments for guide-directed cleavage. We hypothesized that targeting Su9-AmAgo in complex with guide RNAs to this region would result in mtDNA breaks and reduced mtDNA copy number.

**Figure 2.**
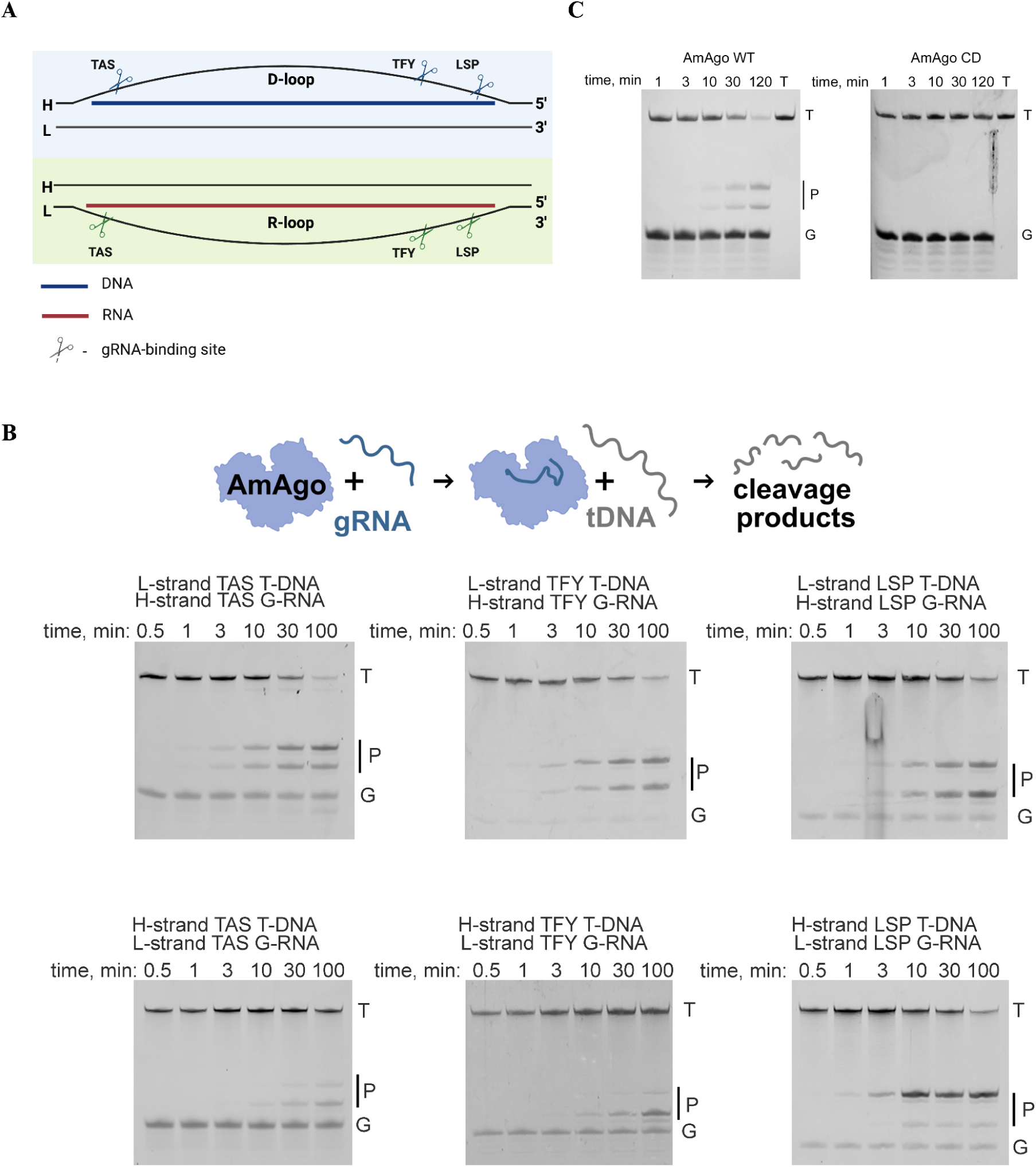
*In vitro* сleavage of guide RNA-directed DNA cleavage by AmAgo. A – **Schematic representation of guide RNA binding sites within the mtDNA control region.** The diagram illustrates the D-loop (top) and R-loop (bottom) single-stranded regions of human mitochondrial DNA, highlighting three selected gRNA target sites within each region – TAS, TFY, and LSP. These sites were chosen for their location near regulatory elements involved in mtDNA replication initiation. Scissor symbols indicate the approximate gRNA binding positions used for targeted cleavage. **B** – **Analysis of target DNA cleavage products generated by wild-type AmAgo.** Recombinant AmAgo protein loaded with indicated gRNAs at 37 °C. Target DNA represents a chemically synthesized ssDNA target, 50 nucleotides in length. Cleavage occurs precisely between the 10^th^ and 11^th^ nucleotides of the guide sequence, counting from the 5’-end. Reaction time points: 0.5, 1, 3, 10, 30, and 100 min. Positions of initial target (T), guide nucleic acids (G), and cleavage products (P) are indicated. **C** – **Analysis of target DNA cleavage products generated by wild-type AmAgo and catalytically inactive AmAgo.**

We selected three loci within the D-loop and R-loop based on their reported single-stranded character and functional significance: the Termination-Associated Sequence (TAS), the Transcription Factor Y-binding site (TFY), and the Light Strand Promoter (LSP) (Figure 2A). For each locus, we designed 18-nucleotide guide RNAs, corresponding to each strand of mtDNA, yielding six distinct guides in total. All guides carried a 5′-terminal adenine, which is essential for AmAgo-mediated cleavage^8^. Synthetic 50-nucleotide ssDNA oligonucleotides corresponding to each target region were used as substrates for *in vitro* cleavage assays.

Incubation of recombinant wild-type AmAgo (AmAgo-WT) with guide RNAs and corresponding target ssDNAs resulted in robust and site-specific DNA cleavage between the 10th and 11th nucleotides relative to the guide 5′ end (scheme in Figure 2B). The expected 23- and 27-nucleotide products were detected by denaturing polyacrylamide gel electrophoresis (Figure 2C), confirming precise endonucleolytic activity. As a negative control, a catalytically inactive AmAgo mutant (AmAgo-CD) with the D331A and D398A substitutions in the PIWI domain exhibited no detectable cleavage at the tested site (Figure 2C).

### AmAgo nuclease programmed with gRNA decreases mtDNA copy number

To evaluate the ability of AmAgo to deplete mitochondrial DNA in living cells, we transduced HEK293T cells with a lentiviral construct encoding Su9-tagged AmAgo or AmAgo without MTS as a control. Forty-eight hours after transduction, synthetic guide RNAs targeting the mitochondrial D-loop and R-loop were delivered via lipid-based transfection. Quantitative PCR analysis revealed an approximately threefold reduction in the mtDNA copy number relative to untreated controls (Figure 3A). All designed guide RNAs induced similar levels of mtDNA depletion, and no statistically significant differences in the efficacy were observed between them. No mtDNA reduction was observed when AmAgo lacked a mitochondrial targeting sequence, even in the presence of guide RNAs, confirming that mitochondrial localization is required for mtDNA depletion.

**Figure 3.**
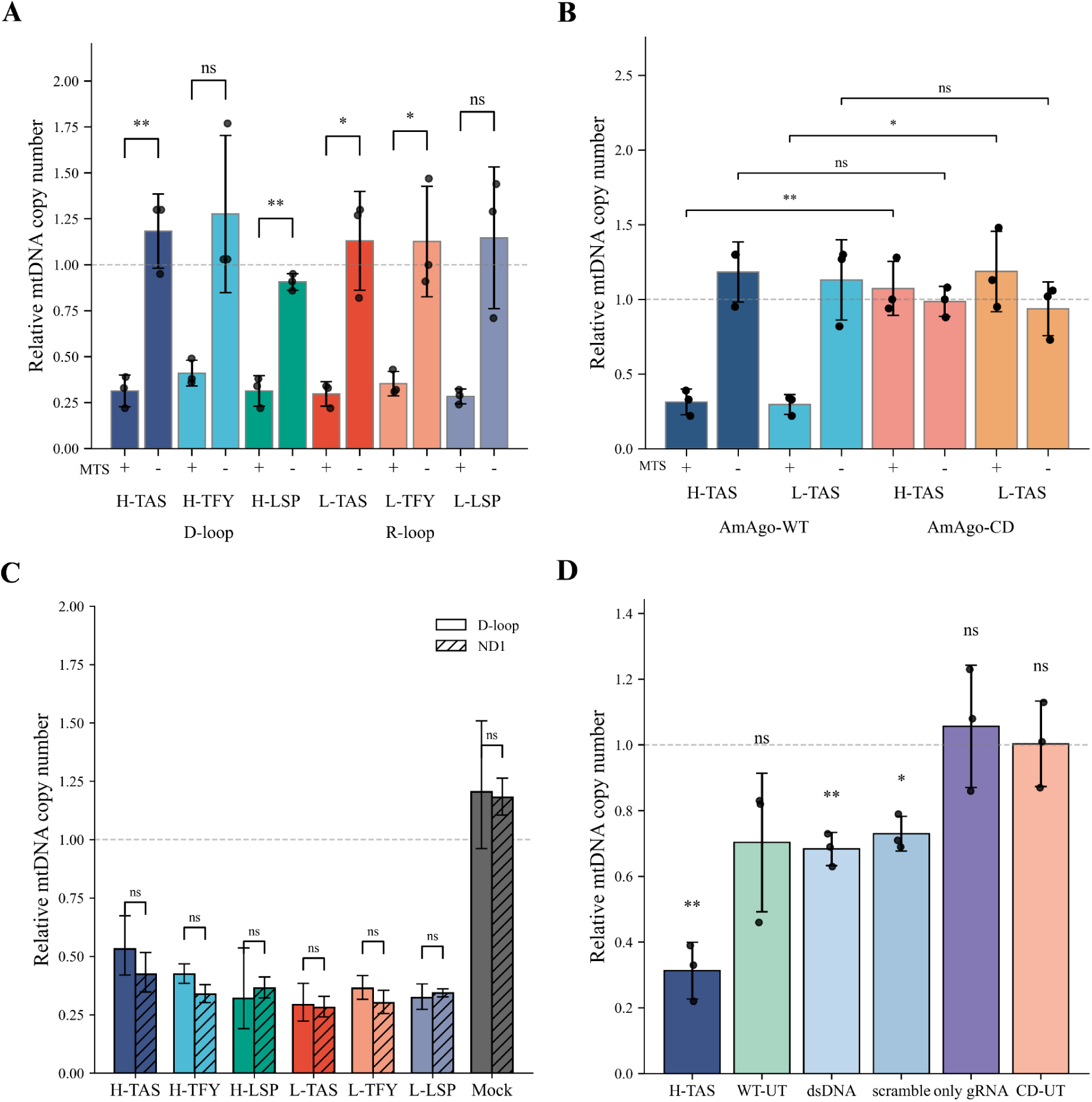
Depletion of mtDNA by wild-type AmAgo. **A** – **Relative mtDNA copy number in HEK293T cells expressing wild-type AmAgo with or without MTS, co-delivered with region-specific guide RNAs.** Left: three guides directed to single-stranded regions of the heavy strand (H-TAS, H-TFY, H-LSP) within the D-loop; right: three guides targeting the light strand (L-TAS, L-TFY, L-LSP) within the R-loop. Expression of AmAgo without an MTS did not alter mtDNA levels. In contrast, AmAgo targeted to mitochondria resulted in a consistent ∼3-fold reduction in mtDNA copy number across all guide designs. Data are represented as mean ± SD (*n* = 3 technical replicates from 3 biological replicates). Statistical significance was assessed using an unpaired two-tailed Student’s *t*-test. **B** – **Relative mtDNA copy number in HEK293T cells expressing wild-type or catalytically inactive AmAgo.** Quantification of mtDNA levels following co-transfection with guide RNAs targeting either the H-strand (H-TAS) or L-strand (L-TAS) of the mtDNA control region. Cells expressed either wild-type AmAgo (AmAgo-WT) or a catalytically inactive mutant (AmAgo-CD), each with or without MTS. AmAgo-WT with MTS and specific guides resulted in a marked reduction in mtDNA copy number (∼3-fold), whereas the catalytically inactive AmAgo-CD had no effect on mtDNA levels, even in the presence of MTS. Data are represented as mean ± SD (*n* = 3 technical replicates from 3 biological replicates). Statistical significance was assessed using an unpaired two-tailed Student’s *t*-test. **C** – **Relative mtDNA copy number in HEK293T cells expressing wild-type AmAgo with MTS and target-specific guide RNAs.** To assess whether mtDNA depletion was restricted to the targeted D-loop region or indicative of global genome degradation, qPCR was performed using primer pairs specific to the D-loop or ND1 gene. Data represent relative mtDNA copy number normalized to nuclear B2M and presented for each guide RNA group (H-TAS, H-TFY, H-LSP, L-TAS, L-TFY, L-LSP). A mock control (Lipofectamine-only transfection) was included for reference. No significant differences were observed between D-loop and ND1 measurements within each group (paired *t*-test, ns), suggesting uniform mtDNA depletion rather than region-specific loss. Data are represented as mean ± SD (*n* = 3 technical replicates from 1 biological replicates). Statistical significance was assessed using a paired two-tailed Student’s *t*-test. **D** – **Relative mtDNA copy number in HEK293T cells transfected with control constructs.** WT-UT and CD-UT represent wild-type and catalytically inactive AmAgo, respectively, without guide RNAs. Additional controls include WT AmAgo co-delivered with a guide targeting a double-stranded region of mtDNA (dsDNA) or a non-complementary scrambled guide (scramble), as well as guide RNA alone (only gRNA). Data are represented as mean ± SD (*n* = 3 technical replicates from 3 biological replicates). One-sample two-tailed *t*-test was performed relative to a hypothetical mean of 1, which corresponds to the average mtDNA copy number in untreated HEK293T cells (HEK293T-UT), used here as a reference baseline for relative quantification. Statistical significance is represented as follows: ns, not significant (P > 0.05); *P ≤ 0.05; **P < 0.01.

In contrast, a catalytically inactive AmAgo mutant failed to affect mtDNA levels (Figure 3B), demonstrating that mtDNA depletion requires an intact nuclease site. This observation also rules out the possibility that mtDNA loss could be due to nonspecific toxicity from protein overexpression, supporting the conclusion that the observed effects are driven specifically by the catalytic activity of wild-type AmAgo.

To determine whether the observed effect reflected global mtDNA depletion rather than localized cleavage at the loop regions, we quantified mtDNA copy number using two independent primer sets targeting the D-loop and the ND1 coding region. Both assays yielded comparable reductions in mtDNA abundance (*p* = 0.180), supporting the conclusion that genome-wide mtDNA degradation had occurred (Figure 3C).

To evaluate potential off-target effects and assess baseline variability in mtDNA levels, we quantified mtDNA copy number in cells expressing wild-type AmAgo in the absence of guide RNAs (AmAgo WT, untreated; WT-UT) (Figure 3D). Although these samples did not exhibit a statistically significant reduction in mtDNA abundance compared to untreated controls (*p* = 0.135), they showed increased variability across replicates (Figure 3D), suggesting the possibility of stochastic loading of endogenous mitochondrial RNAs as unintended guides. In contrast, expression of a catalytically inactive AmAgo mutant in the absence of guides (AmAgo-CD, untreated; CD-UT) had no measurable effect on mtDNA levels (*p* = 0.969), indicating that the observed fluctuations in the WT-UT condition depend on nuclease activity. Likewise, transfection of guide RNAs alone, without AmAgo protein, did not affect the mtDNA copy number (*p* = 0.650).

Finally, to investigate whether specific targeting of the D-loop and R-loop regions by guide RNAs is important for mtDNA depletion, we tested synthetic guide RNA targeting a predicted double-stranded region of mtDNA corresponding to the MT-RNR2 gene and scramble guide RNA with no complementarity to the mitochondrial genome (Figure 3D). These guides caused a reproducible and modest (∼1.5-fold) reduction in mtDNA levels (*p* = 0.008 and *p* = 0.013, respectively), with less variability compared to the WT-UT condition. Therefore, AmAgo retains some ability to reduce mtDNA copy number even in the absence of exogenous guide RNAs targeting mtDNA. However, these effects are reproducibly lower compared to the conditions with guide RNAs targeting the R-loop and D-loop regions.

## Discussion

In this study, we present the first targeted manipulation of human mtDNA using a pAgo protein, AmAgo. By fusing AmAgo to a mitochondrial targeting sequence and co-delivering synthetic guide RNAs complementary to the mitochondrial D-loop or R-loop regions, we achieved a robust and reproducible reduction - approximately threefold - in mtDNA copy number within 48 hours. These results suggest that pAgo-based tools could serve as a programmable approach to modulate mitochondrial DNA copy number.

To test the hypothesis that AmAgo remains functionally active upon mitochondrial import, we selected the mitochondrial control region - a key locus for the initiation and regulation of mtDNA replication comprising the D-loop and R-loop structures - as a target.^25^ This region was chosen due to its well-characterised single-stranded features, which are essential for mtDNA replication and are particularly well-suited for cleavage by pAgos, such as AmAgo, which operate on single-stranded DNA substrates.^7,8,10,26,27^ The D-loop contains a short third DNA strand (7S DNA) annealed to the heavy strand, forming a triple-stranded region that regulates replication initiation and maintenance. The adjacent R-loop includes RNA-DNA hybrids formed by long non-coding RNAs (LC-RNAs) transcribed from the heavy-strand promoter, playing pivotal roles in initiating and regulating mtDNA replication, nucleoid stability, and DNA segregation.^25^ The partially unpaired and accessible nature of the D- and R-loop structures within the mitochondrial non-coding control region provided an opportunity to assess whether RNA-guided AmAgo could induce site-specific cleavage and reduce mtDNA copy number.

The nearly threefold reduction (70.8 ± 4.0%) in mtDNA copy number observed upon delivery of AmAgo in complex with synthetic guide RNAs is consistent with a disruption of essential mitochondrial regulatory mechanisms. Cleavage within the D-loop or R-loop likely compromises structural elements required for the initiation of strand-asynchronous replication, thereby hindering the formation of functional replication intermediates.^25,28^ In addition, such targeted cleavage may interfere with the association of mtDNA with the inner mitochondrial membrane, which is critical for nucleoid organisation and genome segregation.^29^ Collectively, these effects could destabilise mitochondrial genome maintenance, leading to a reduction in replication events and the observed decline in mtDNA copy number.

To confirm that the observed effects depended specifically on mitochondrial localisation and the catalytic activity of AmAgo, we performed a series of control experiments. AmAgo lacking the mitochondrial targeting sequence was localised to the cytoplasm and nucleus and had no effect on mtDNA copy number, indicating that mitochondrial import is essential for its activity. Similarly, a catalytically inactive AmAgo mutant did not alter mtDNA copies. These results demonstrate that mtDNA depletion requires both mitochondrial localisation and nuclease activity, and is mediated specifically by guide RNA-directed cleavage within mitochondria.

In addition, we observed a modest but reproducible reduction in mtDNA copy number upon mitochondrial delivery of wild-type AmAgo in the absence of synthetic guide RNAs. While this effect did not reach statistical significance (*p* = 0.135), the increased variability across replicates suggests that endogenous mitochondrial RNAs may serve as unintended guides. Similarly, modest but reproducible decrease in the mtDNA content was also observed in the presence of synthetic guide RNAs corresponding to a dsDNA region or noncomplementary to mtDNA. The human mitochondrial transcriptome contains a variety of RNA species, including tRNAs, processed mRNAs, rRNA fragments, and small non-coding RNAs, many of which possess 5′-adenosine termini and lengths compatible with prokaryotic Argonaute loading.^30,31,32^ Some of these RNAs are mitochondrially encoded, while others are imported from the cytosol.^33,34^ Even rare stochastic engagement with accessible single-stranded regions such as the D-loop may suffice to trigger mtDNA destabilization. This interpretation is supported by the control experiments, since catalytically inactive AmAgo without synthetic guide RNAs had no measurable effect on mtDNA levels, and cells transfected with guide RNAs alone also showed no significant change.

Together, these results suggest that rational guide selection and structural accessibility of the target site can influence the efficiency and specificity of mtDNA depletion. These findings highlight the potential for low-level off-target activity driven by endogenous RNAs, an issue shared with CRISPR-Cas systems when they were just starting to optimize.^35,36^ Therefore, mapping the repertoire of endogenous mitochondrial RNAs and refining guide selection will be critical to enhancing the specificity and safety of pAgo-based editing tools.

Argonaute proteins offer several distinct advantages over CRISPR-Cas systems, including their smaller protein size, shorter guide nucleic acid requirements, and independence from DNA motifs such as PAM sequences.^4,5^ These attributes not only facilitate more efficient mitochondrial import but also expand editing capabilities beyond those achievable with traditional mitochondrial genome editing tools. Previous studies have highlighted challenges associated with mitochondrial import of RNA that are often constrained by RNA length and structural complexity.^37^ Our recent research indicates that shorter RNA molecules, comparable in length to Argonaute guides, effectively enter mitochondria without additional import determinants,^38^ suggesting intrinsic mitochondrial targeting advantages inherent in Argonaute-guided RNAs. Furthermore, while other programmable nuclease systems have been associated with cytotoxicity and unintended effects on mtDNA maintenance in some contexts,^35^ cells expressing AmAgo showed no signs of toxicity and remained viable during long-term cultivation, with no observable morphological abnormalities. These observations support the overall biocompatibility of the AmAgo system and its potential suitability for applications requiring sustained mitochondrial manipulation.

Beyond simply reducing mtDNA copy numbers, Argonaute proteins hold potential for advanced mitochondrial genome editing applications, including precise site-specific deletions (such as targeting single-stranded DNA of the major arc during mtDNA replication^39^) or targeted base editing through fusion with deaminase domains.^40^ The broad implications of our findings suggest therapeutic possibilities for mitochondrial diseases. Future research efforts should focus on enhancement of guide RNA stability and specificity, and refinement of mitochondrial editing capabilities. Successful development of Argonaute-based mitochondrial genome editing technologies would significantly advance our mitochondrial genome engineering toolbox, fostering progress in fundamental mitochondrial biology research and clinical therapeutic interventions.

## Acknowledgments

We thank Anna Demchenko and Irina Panchuk (Research Centre for Medical Genetics) for assistance with flow cytometry experiments.

## Author contributions

Conceptualization, I.M., A.K.; Investigation, B.R., E.S., E.K., L.L., D.G., E.U.; Resources, E.U., V.S., S.S.; Formal Analysis, B.R.; Visualization, B.R., L.L.; Writing – Original Draft, B.R.; Writing – Review & Editing, E.K., L.L., D.G., S.S., A.K., I.M.; Supervision, I.M., A.K.

## Supplemental information

Document S1. Includes Tables S1–S3.

